# NanoBEP – A Machine Learning Based Tool for Nanobody Binding Energy Prediction

**DOI:** 10.1101/2025.02.04.635413

**Authors:** Soma Prasad Sahoo, Pratibha Manickavasagam, M. Hamsa Priya

## Abstract

Nanobody is a special class of antibodies comprising only one variable heavy chain. Its small size and high stability over a wide range of temperature and pH, makes it an ideal candidate for biomedical applications. Designing a nanobody that can bind to a specific target protein, either for therapeutic or diagnostic purposes, requires a quick estimation of binding affinity of nanobody-protein complex. Many predictive models for protein-protein interactions have been developed leveraging the capability of machine learning techniques. The popular protein-protein interaction models, however, could not accurately predict the binding affinity of available nanobody-protein complexes. We, therefore, have developed a random forest based model that can predict the value of dissociation constant (log_10_ K_d_) at high accuracy with a Pearson’s correlation coefficient value of 0.95 and a mean absolute error of 0.44. Our cherry-picked model identifies the best protein features for the prediction through two stages of selection strategy that includes elimination of highly correlated features through graph network analysis, followed by the recursive feature elimination through random forest. Despite being a class of antibodies, a model trained only on antigen-antibody complexes couldn’t accurately predict the binding affinity of the nanobody-protein complexes. The predictability improved only when we included the data on monomeric protein complexes and some nanobody-protein complexes during training.

## 1 Introduction

Nanobodies are a novel class of antibodies, currently gaining importance owing to its small size, high stability and a wide range of applications in therapeutics and diagnostics [1]. Structurally, an antibody contains two identical heavy chains and two identical light chains with each chain comprising of a variable and a constant regions. On the other hand, a nanobody contains only one variable heavy chain, otherwise referred as variable heavy chain only antibody (VHH), making them weigh only about 1/10^th^ of that of an antibody. Interestingly, nanbodies are highly thermo stable maintaining their structure even upto a temperature of 95°C and can withstand a wide pH range [1, 2]. The stable interaction among the four beta sheets corresponding to the framework regions (FRs) including a disulphide bond attribute to the structural stability of the nanobody. [3]

Engineering the disordered complementarity determining regions (CDRs) to enable strong binding to the antigen or any specific protein as the target helps in developing a desired therapeutic or diagnostic agent. Nanobodies based FDA approved drugs are available for cancer and rheumatoid arthritis [4] and potential drug candidates have been identified for treating tropical diseases such as dengue, rabies and so on [5]. The conventional route to identify target specific nanobody is through immunizing the animals belonging to the family Camelidae, followed by a tedious isolation and purification processes [1]. An alternative approch is through PHAGE DISPLAY technique with subsequent laborious screening for the desired binding strength [6].

The dissociation constant (*K*_*d*_) is the most common thermodynamic parameter characterizing the strength of binding of a protein complex [7]. Typically, *K*_*d*_ values can be in the range of micromolar to picomolar depending on whether the protein-protein interaction is either transient or permanent association [8]. Some of the experimental techniques for determining *K*_*d*_ values include yeast two-hybrid (Y2H) assays [9], Fluorescence Resonance Energy Transfer (FRET) [10], Surface Plasmon Resonance (SPR) [11], and Isothermal Titration Calorimetry (ITC) [12]. On the other hand, several computational models based on the existing experimental data, for predicting *K*_*d*_ values have also been developed to cut down the experimental cost. Such models provide a quick screening strategy to eliminate weak binders during protein design.

Over the past decade, various web servers for predicting the binding affinity of protein complexes through either sequence [13] or structure [14] based features have been developed. These web servers were built using machine learning (ML) techniques; the recent ones leveraging the advancement in computational techniques and hardware usually provide a rapid and accurate prediction. The underlying ML models can be classified as linear models (e.g., linear regression, support vector machines, regularized linear models), nonlinear models (e.g., decision trees, random forest, neural networks), and mixed models (e.g., hierarchical bayesian models, linear mixed-effects models) [15, 16]. For instance, Yugendar et al., devised a linear regression model based on protein functional classification to forecast binding affinity from sequence based parameters and the performance of the model in terms of Pearson’s correlation coefficient (R value) is about 0.73–0.99 as evaluated by Jack-Knife test [17]. The most popular PROtein binDIng enerGY (PRODIGY) server employing linear regression with features computed from both the interacting and non-interacting residues in three-dimensional structures exhibits a R value of 0.73 [18, 19].

Through a graph-based machine learning approach, the web server CSM AB could predict antigen-antibody binding affinity with a R value of 0.64 [20]. Another web server, PPI Affinity, developed by Sandra et al., employs support vector machine and achieves a R value of 0.63 for predicting binding energy in protein complexes [21]. *AREA Affinity* web server devised by Yang et al., on the other hand, considers interface and surface area based features for predicting protein-protein binding energy using a combination of linear, non-linear algorithms (neural network and random forest) and mixed-models (linear combination of nonlinear models) and their best model was the area-based neural network model exhibiting a R value of 0.87 [16]. DeepPPAPred web server leveraging deep learning technique on a diverse array of features extracted from protein sequences and structures, predicts antigen-antibody binding affinity with a R value of 0.79 [22].

We believed that the design of nanobodies can be accelerated with the available advanced protein-protein interaction models. The prediction of binding affinity of known nanobody-protein complexes, however, through the web servers discussed above was extremely poor. This prompted us to develop a new ML model to predict *K*_*d*_ values for nanobody-protein complexes. We initially tried to construct a support vector regression (SVR) model, but the performance was only moderate, therefore, we opted for a random forest (RF) model to enhance the performance. Our model was trained using available antigen-antibody structures, monomeric protein complexes and some nanobody-protein complexes. Although, nanobody is a subclass of antibody, the model performance was good when we considered monomeric protein complexes alone or along with the antigen-antibody complexes, since the former included more experimental data over a wide range of *K*_*d*_ values. The predictability of RF model is found to be excellent only when we included some nanobody-protein complexes during training. Our best RF model exhibits a R value of 0.95 and a mean absolute error of 0.44; the robustness of the model was further validated through bootstrap sampling.

## 2 Methods

### 2.1 Data Collection

We gathered experimental binding energy data for 76 nanobody-protein (Nb-P) complexes [20], 342 antigen-antibody (Ag-Ab) complexes [20, 23] and 524 monomeric proteinprotein (P-P) complexes [24]. We ensured that all these 942 complexes are non-redundant and have also cross-verified their dissociation constants (K_*d*_) to the values listed in PDB-bind (v.2020) [25] database. We, however, found mismatch in binding free energy, Δ*G* ≡ *RT* ln *K*_*d*_, for some protein complexes between the servers, and realized that it is because of the difference in temperature used to convert *K*_*d*_ to Δ*G* and *vice versa*, hence, we meticulously garnered the temperature at which the binding energy was characterized for each of the complexes from the corresponding research article. The binding disassociation constant, *K*_*d*_, for the protein complexes in the acquired dataset ranges from 10^−3^ to 10^−15^ M. Considering the variation in the several orders of magnitude, we used log_10_ K_*d*_ values to train our model.

### 2.2 Feature Generation

A total of 134 features were employed in our model. As temperature is an important thermodynamic parameter and there exists some variation in the temperature among the binding energy characterization of protein complexes, we used temperature as a feature in our model. The remaining 133 features include both the sequence and structure based descriptors. We have excluded all the HETATOMS from the PDB file to compute the structure based features. Table. S1 provides the list of all the features with a detailed description about them. We extracted eight features describing the type of amino acids – polar (H, N, C, Q, W, T, S), non-polar (G, A, F, I, V, L, P, M, Y) and charged (R, K, D, E) amino acid residues – that are in contact at the the interface of the protein-complex and on their non-interacting surfaces (NIS) from https://github.com/haddocking/prodigy [18]. Next eight features corresponding to the solvent accessible surface area (SASA) *i*.*e*., receptor surface area (RSA) and ligand surface area (LSA) were extracted from https://github.com/nioroso-x3/dr\_sasa\_n [26]. Both RSA and LSA were also classified based on the amino acid type – basic, acidic, non-polar, polar.

Further, we have included 77 interaction energy and entropy features computed via protein-protein Δx (ppdx) program [27] (https://github.com/simonecnt/ppdx). This includes features derived from fourteen different software packages – CHARMM (24), MD-Traj (4), Native MDTRaj (9), Biopython (7), pyDock (4), RFPP(2), iPot(4), Modeller (5),Rosetta (3), Numpy (2), ATTRACT (1), RFPP, ZRANK(2), FoldX (9). The number in the brackets indicates the number of features obtained from the respective package. Finally, we employed amino acid pair-wise potentials (AAPPs) to evaluate the strength of interaction between the amino acids that are in contact at the protein interface. We identified a pair of amino acid residues on either proteins in the protein-complex to be in contact if the distance between any of the heavy atoms of the corresponding residues is within 5 Å. The summation of AAPP value for every interacting pair provides the net contact potential (Figure 1A). We used each of the forty available AAPPs [28, 29] to compute a feature, contributing to the last 40 features. In the case of a antigen-antibody (Ag-Ab) complex, all the interacting pairs between the antigen and both the heavy chains (H) and the light chains (L) of the antibody were considered together as shown in Figure 1B.

**Figure 1:**
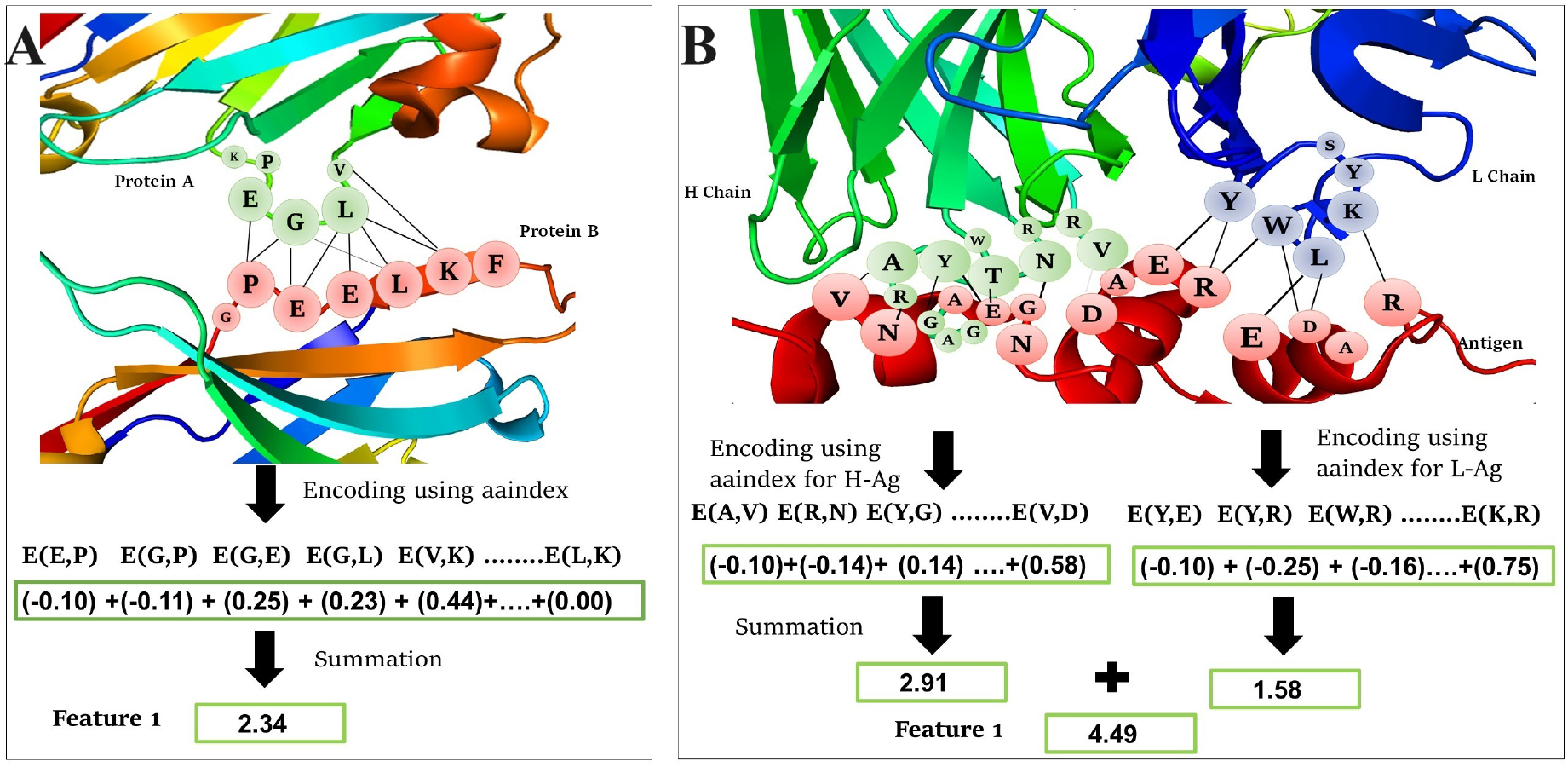
Pictorial representation of encoding protein contact strength using amino acid pairwise potentials (AAPP) as a feature for (A) monomeric protein-protein and nanobody-protein complexes, (B) antigen-antibody complex, involving interaction of antigen (Ag) with both the heavy (H) chains and the light (L) chains of the antibody. E(X, Y) denotes the amino acid pairwise potential between interacting amino acids X and Y. The black lines connect the interacting amino acids between the proteins.

### 2.3 Elimination of Highly Correlated Features

To improve the accuracy in the prediction using the ML model, it is essential to have an optimal non-redundant set of features. We, therefore, employed the NETCORE (network-based, correlation-driven redundancy elimination) algorithm [30] to eliminate the redundant, especially the highly correlated features. A brief description of the algorithm is as follows – first, the correlation coefficient matrix is constructed from the Pearson’s correlation coefficient (R) for every pair of features. Next, a graph was constructed with features as nodes as in Figure 2A and the edges are drawn between those nodes when their corresponding features are highly correlated *i*.*e*., |*R*| > 0.85. The isolated/unconnected nodes represent the first set of uncorrelated or weakly correlated features.

**Figure 2:**
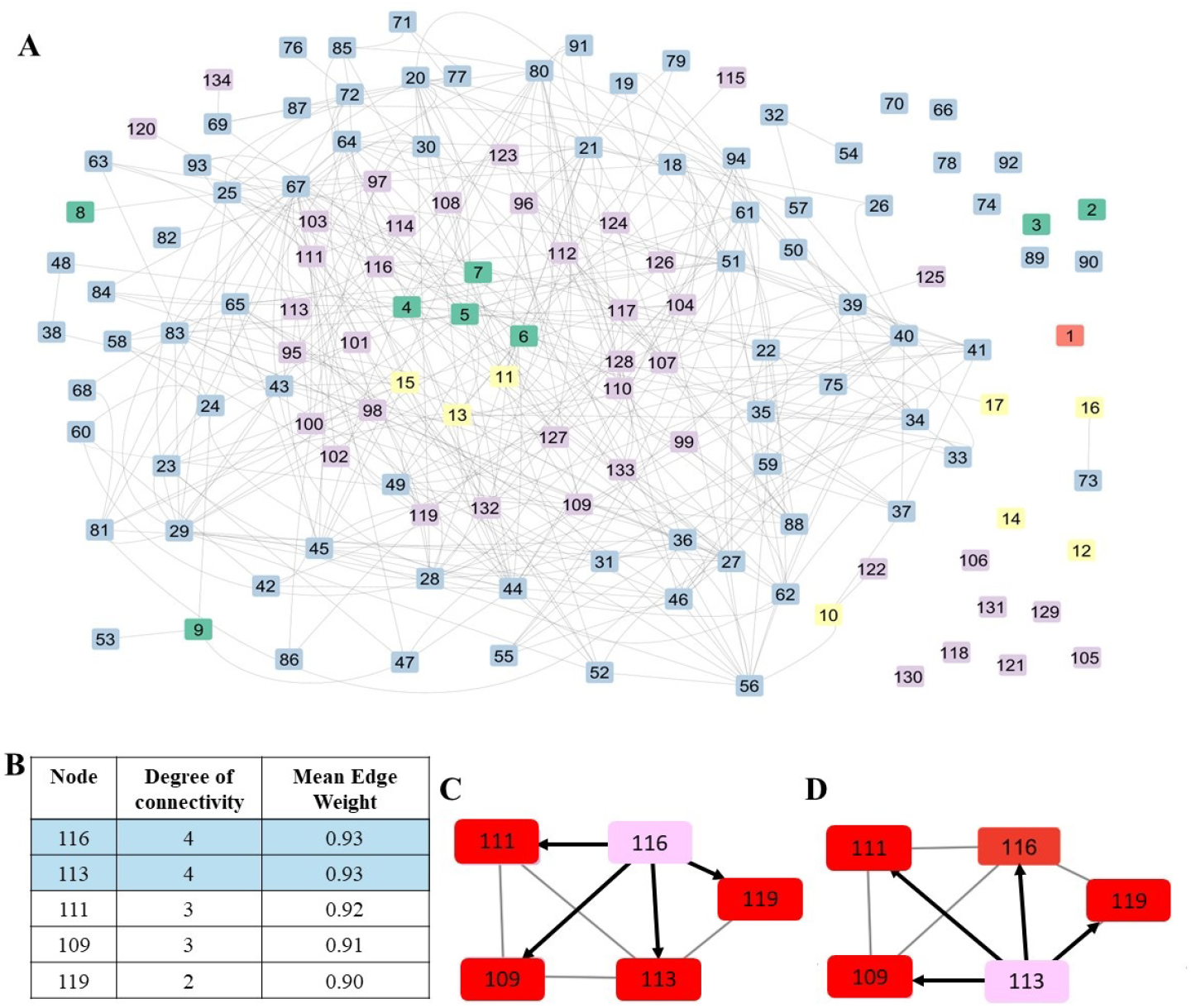
Eliminating highly correlated features through network based scheme: (A) A graph network representation of all the features (nodes) for P-P data. Color code: red box – temperature; green boxes - 8 PRODIGY contact features; yellow boxes – 8 SASA based features; blue boxes – 77 ppdx features; purple boxes – 40 AAPP features; grey lines - edges connecting highly correlated features. (B–D) Illustration of selection of important feature based on the degree of connectivity and mean edge weight by eliminating the other connected nodes.

Then among the connected nodes, the degree of connectivity of each node is calculated.

The node with the highest connectivity is retained as important node in the network, while the nodes connected to them are eliminated. This process is carried out iteratively till we obtain a completely unconnected graph. During the iterative scheme, there might arise a situation that multiple nodes have the same degree of connectivity, in such a scenario, the node with the highest mean edge weight (MEW) is retained and the nodes connected to that node are eliminated. In case of multiple nodes with same degree and mean MEW, we randomly chose one of them as the important node, however, we have verified that our random choice didn’t alter the list of optimal features. In Figure 2 B, both the nodes 113 and 116 have the same degree of connectivity and MEW. As shown in Figure 2 C–D, our choice to retain either the node 116 or node 113, results in the elimination of the other connected nodes like 109, 111 and 119. We used Cytoscape [31] for the network construction and degree analysis.

### 2.4 Modeling Protocol

The dataset comprising crystal structures of 942 protein complexes was divided into three sets (A, B and C) for training and testing our ML model (Figure 3). Set A includes all the protein complexes, with classical 80:20 split ratio for training and testing the ML model, respectively. In set B, ML model was trained using Ag-Ab and P-P complexes and the performance of the model to predict the binding affinity for Nb-P complexes was tested. This set helps to compare the performance of our model with the existing protein-protein binding energy prediction models, since the latter were not modelled using data on Nb-P complexes. In set C, we included all data of Ag-Ab and P-P complexes and 80% of Nb-P complex for ML training, while 20% of the Nb-P data was used as the test set. This third set helps us to comprehend the importance of employing available, though limited, data on Nb-P complexes in training and assess its accuracy of prediction for unseen Nb-P data. Table. S2 list the subsets of these three datasets along with size of the data for training and testing the ML models.

**Figure 3:**
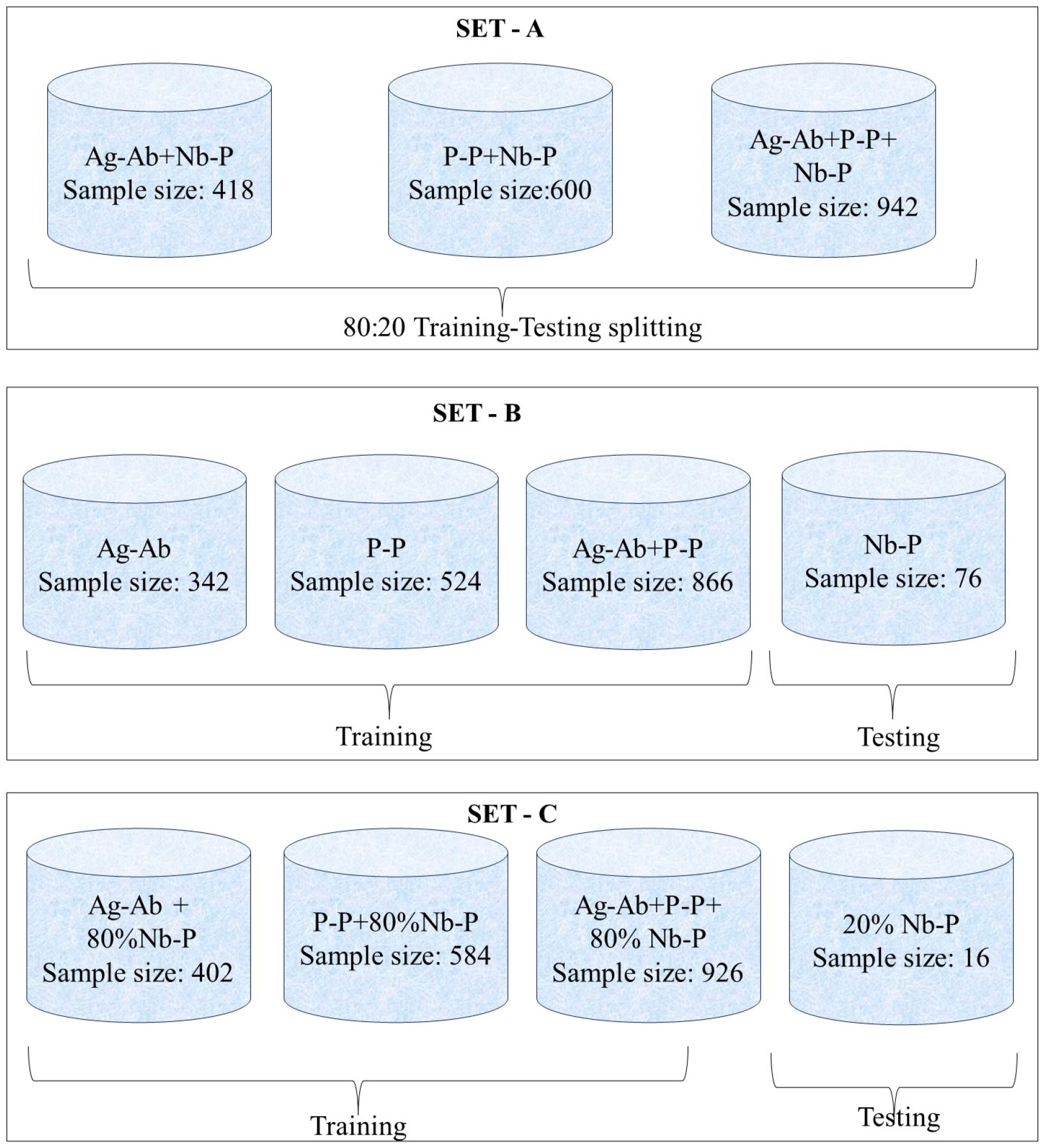
Scheme of data sampling employed for training and testing models

First, we tried a simple ML model through linear support vector regression (SVR) to predict log_10_ *K*_*d*_ of the protein complexes. SVR is an effective supervised learning technique for small to medium sized datasets. It includes two model hyperparameters – epsilon, *ϵ*, setting the tolerance margin for errors and a regularization parameter, C, controlling the trade-off between maximizing the margin and minimizing the error in prediction. We obtained two variants of SVR models for the sets A and B. In the first variant, all 134 features were used and in the second variant only the uncorrelated features obtained from NETCORE algorithm were used.

Next, we developed a random forest model, an ensemble machine learning model that operates on the principle of bootstrap aggregation *i*.*e*., bagging. The model involves construction of multiple decision trees during training, utilizing random subsets of training data and features [32]. Further, we employed the recursive feature elimination technique that iteratively eliminates less significant features, to identify the most critical features.

Through this technique, we could identify the top 20 important features from the set of uncorrelated features obtained by NETCORE algorithm. Subsequently, we developed a procedure to evaluate various possible combinations of these top 20 features to determine the optimal set of 10 important features. Through multiple iterations for randomly selecting the features, the top 10 influential features were determined in a robust manner that improves the model accuracy. Hyperparameters of the model includes the number of trees in the forest (n estimators), maximum tree depth (max depth), minimum samples required at a leaf node (min samples leaf), and the random state.

We used Scikit-learn package [33] to obtain both SVR and RF models. The model efficacy was assessed using Pearson’s correlation coefficient, 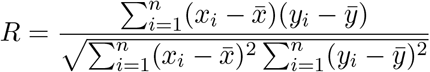 where *x*_*i*_ and *y*_*i*_ correspond to the actual and predicted values, respectively, and the 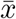 and 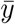 imply their corresponding mean values for *n* observations, and by the mean absolute error,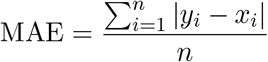. The hyperparameters of the models were optimized through grid search technique over the parameter space using 10-fold cross-validation to obtain the best R value with the lowest MAE. We generated multiple models and identified the random state that yielded the highest accuracy [27], in other words, effectively “cherry-picking” the best model. This is indeed a two stage process, in the first stage we identified the best random state for train and test data split over 50,000 iterations, in the second stage we once again carried out 50,000 iterations to determine the best random state for the model. The histograms of correlation coefficients obtained in the process for all data sampling schemes are shown in Figure S3. The models corresponding to the mean (average R value) and the right most data (highest R value) on the histogram are referred as the average and the best cherry-picked models, respectively. The cherry picked model represents the maximum performance with the best combination of features and hyper-parameters. Further, the robustness of the chosen models are checked through bootstrap sampling on test data. The histograms of correlation coefficients during bootstrapping for the average and the cherry-picked models shown in figure S4, provides a visual comparison of dependability and variability of the models.

## 3 Results and Discussions

### 3.1 Need to Develop a Nanobody Binding Energy Prediction Model

The ability to design nanobodies targeting a specific receptor might pave way for effective therapeutics/diagnostics and also for personalised medicine. For a systematic designing, it is essential to predict the binding affinity of nanobody to the receptor accurately. We utilized several web servers, *viz* PRODIGY, *Area Affinity*, DeepPPAPred, CSM AB, and PPI Affinity, to predict the *K*_*d*_ values for the 76 Nb-P complexes we have collected. All these web servers require only the PDB structure as an input for their prediction. Table 1 list the underlying ML model of the server along with the R and MAE values for the predicted *K*_*d*_ values as against their experimental values for the Nb-P complexes. We can observe that the performance of all the servers is poor with a R value close to *zero* for three servers and weak negative correlation with R values of -0.223 and -0.148 for PPI Affinity and CSM AB, respectively; the MAE is greater than 1.5 for all the servers. This prompted us to develop a ML model that can accurately predict binding affinity for Nb-P complexes.

**Table 1:** Performance of popular web servers on protein-protein interactions in predicting *K*_*d*_ values for nanobody-protein complexes.

| Web Server | ML method | R value | MAE |
| --- | --- | --- | --- |
| PRODIGY | Linear Regression | 0.03 | 1.59 |
| *Area Affinity* | Linear and Non-linear methods | 0.03 | 1.50 |
| PPI Affinity | Support Vector Machine | -0.22 | 1.52 |
| CSM AB | Graph based model | -0.15 | 2.30 |
| DeepPPAPred | Deep learning model | -0.01 | 1.52 |

R – Pearson’s correlation coefficient; MAE - mean absolute error

We collected a total of 942 complexes including Ag-Ab, P-P and Nb-P complexes. The *K*_*d*_ values for these complexes vary in the range 10^−3^ to 10^−15^ M. In order to faithfully capture such a large variation spanning over 12 orders of magnitude, we choose to use log_10_ *K*_*d*_ in our model. The top panel of Figure 4 shows the probability distribution of binding affinity (log_10_ *K*_*d*_) values for antigen-antibody, monomeric protein-protein and nanobody-protein complexes. The range of binding affinity values is lower for Ag-Ab and Nb-P complexes with a mean *K*_*d*_ value of 10^−8^–10^−9^ M. As all protein-protein interactions need not be as specific as Ag-Ab interactions, the P-P complexes also includes *K*_*d*_ in the lower range 10^−3^–10^−6^ M, hence their mean is slightly lower about 10^−7^–10^−8^ M.

**Figure 4:**
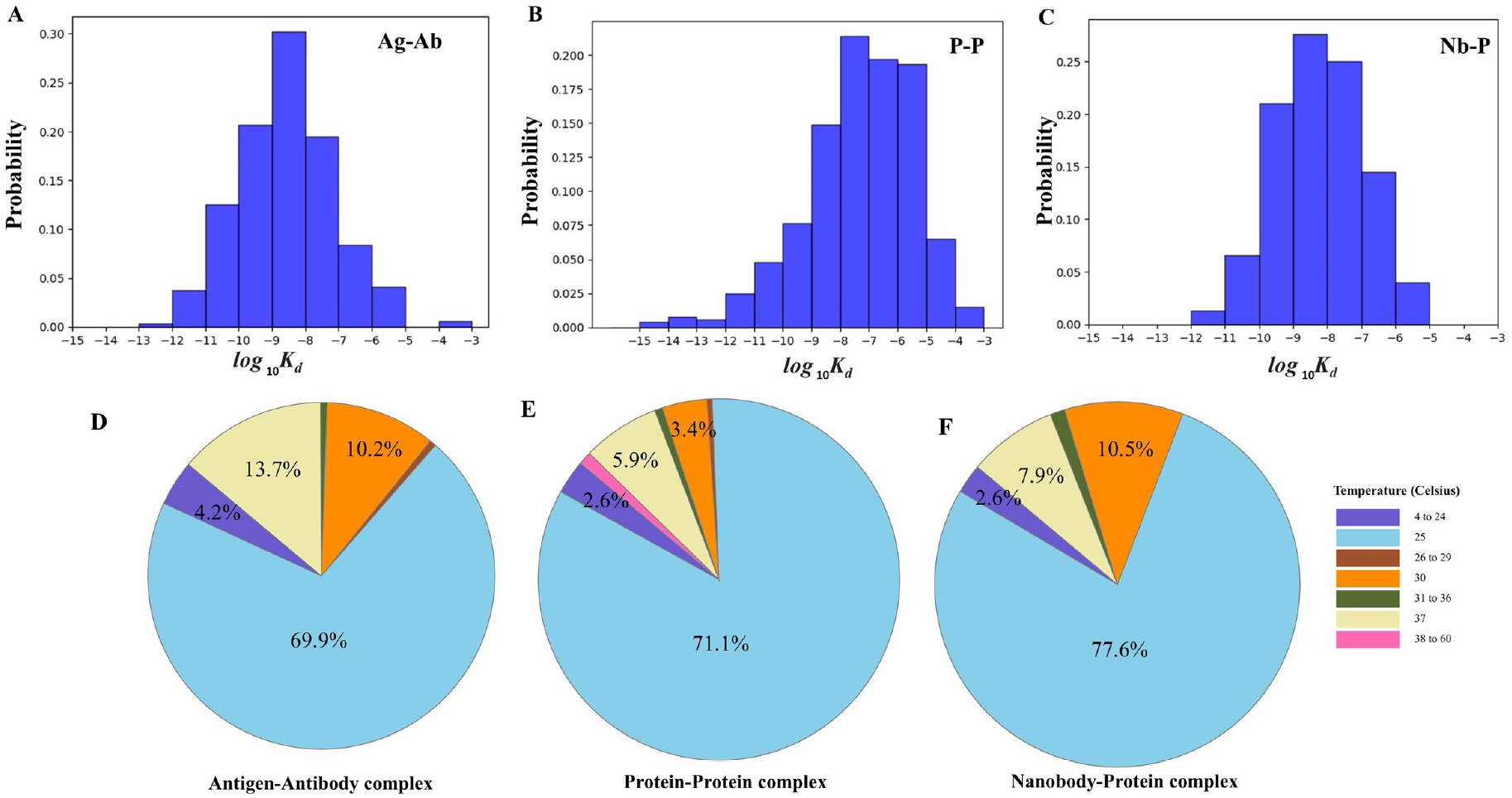
Statistics on the binding affinity (top panel) and the temperature (bottom panel) at which the binding affinity was measured for various protein complexes: A & D – antigen-antibody complexes (Ag-Ab); B & E – protein-protein complexes (P-P); C & F – nanobody-protein complexes (Nb-P)

The bottom panel of Figure 4 shows the pie-chart on the temperature at which the binding affinity was experimentally measured for the protein complexes. For most of the complexes, about 70-78% of the data, *K*_*d*_ had been determined at 25°C, yet for the remaining 30% data, the temperature of measurement was either close to 4°C or to the physiological temperature of 37°C. For example, K_*d*_ values for Ag-Ab and Nb-P complexes were measured at 30°C – 37°C for 24% and 20% of the data, respectively. The binding affinities for only 2-4% of Ag-Ab and Nb-P complexes were characterized at lower temperatures, mostly 4 °C and 15°C, whereas, ∼18% of P-P complexes were characterized at lower temperatures. The *K*_*d*_ measurement of P-P complexes at higher temperatures (30°C – 37°C) was for ∼10% of the data. The observed variation in temperature between the experiments suggested us to consider temperature as an important feature in our model, since temperature is a key thermodynamic parameter determining the energetics of protein binding.

### 3.2 Feature Selection using Network based Elimination Method

Figure 2A represents all 134 features as a graph network. The nodes represent the features and edges are drawn only between highly connected nodes *i*.*e*., |*R*| > 0.85. The color of the nodes represent the different classes of features we have considered as described in Methods section. From the graph network, we can clearly see that only 19 nodes are completely uncorrelated, all other nodes are connected to at least one another node in the network. These 19 isolated nodes correspond to temperature, 2 PRODIGY features, 2 SASA based feature, 7 ppdx and 7 AAPP features. So, only ∼10% of ppdx and 18% of AAPP features are uncorrelated among all features. It is, therefore, redundant to consider all the features and therefore, adopted a strategy to eliminate highly correlated features using NETCORE algorithm [30].

Figure 5A depicts the extent of correlation among the features as a heat map. Here again we can notice two domains (red patches) indicating the high correlation among many of ppdx and AAPP features. After processing the features through NETCORE algorithm (see Methods), we could eliminate ∼55% of features as they are highly correlated. The heat map in Figure 5B shows more blue patches. Some orange spots can be observed as well but they correspond to features that are the moderately correlated (0.6 < |*R*| < 0.85), since we have eliminated only the highly correlated (|*R*| > 0.85) features through NETCORE. It is also to be noted that the set of non-redundant features differs for the different input data such as all the protein complexes or Ag-Ab+Nb-P complexes or monomeric P-P+Nb-P complexes and some other combinations thereof.

**Figure 5:**
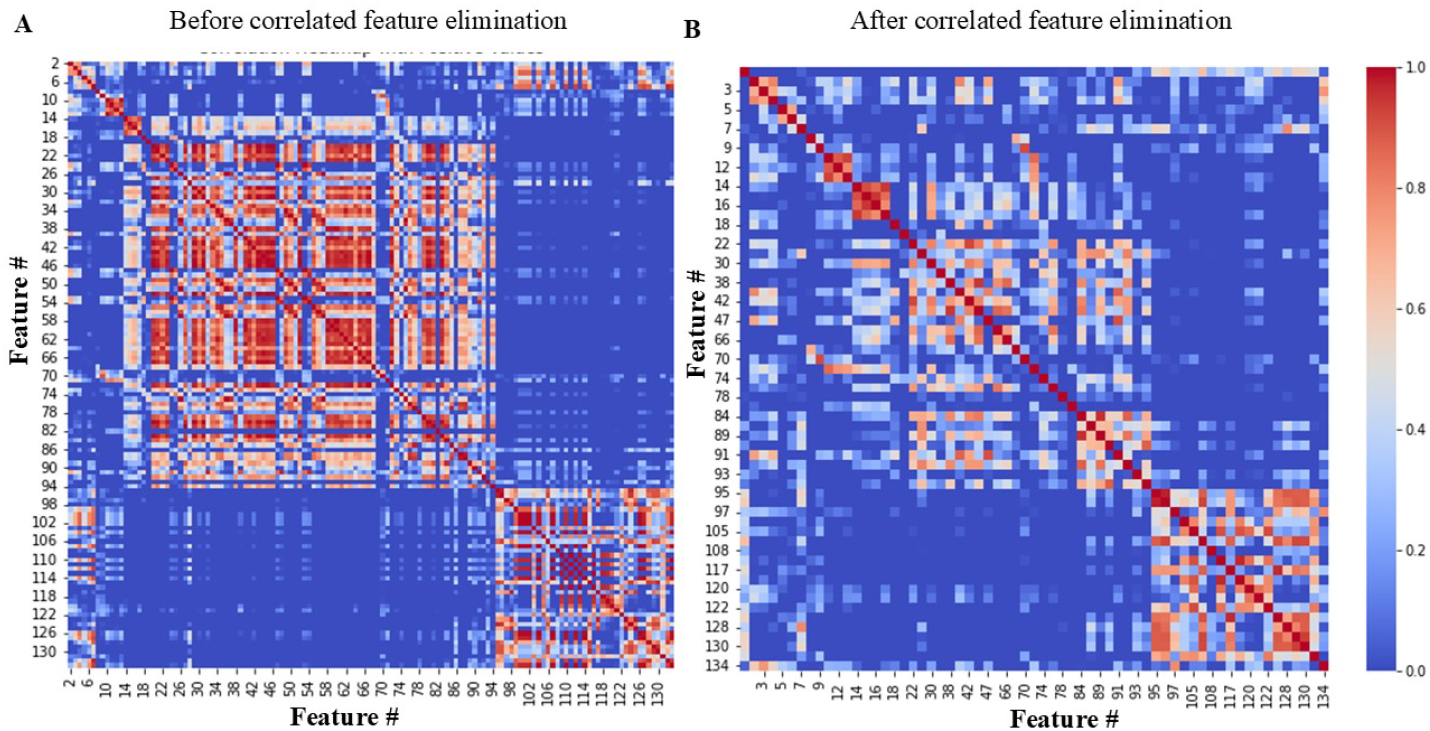
Heat map showing the extent of correlation for P-P data A) among all 134 features and B) after removal of highly correlated features using NETCORE algorithm for the dataset B2 (Table S2).

### 3.3 Performance Analysis of Support Vector Regression (SVR)

With an interest to develop a simplest predictive model, we started with support vector regression (SVR). Figure 6 presents the performance of the SVR model with non-redundant features obtained through NETCORE algorithm. For comparison, we had also obtained SVR model based on all features, the corresponding performance plots are shown in Figure S2. The performance of both the variants of SVR model, however, are almost the same.

**Figure 6:**
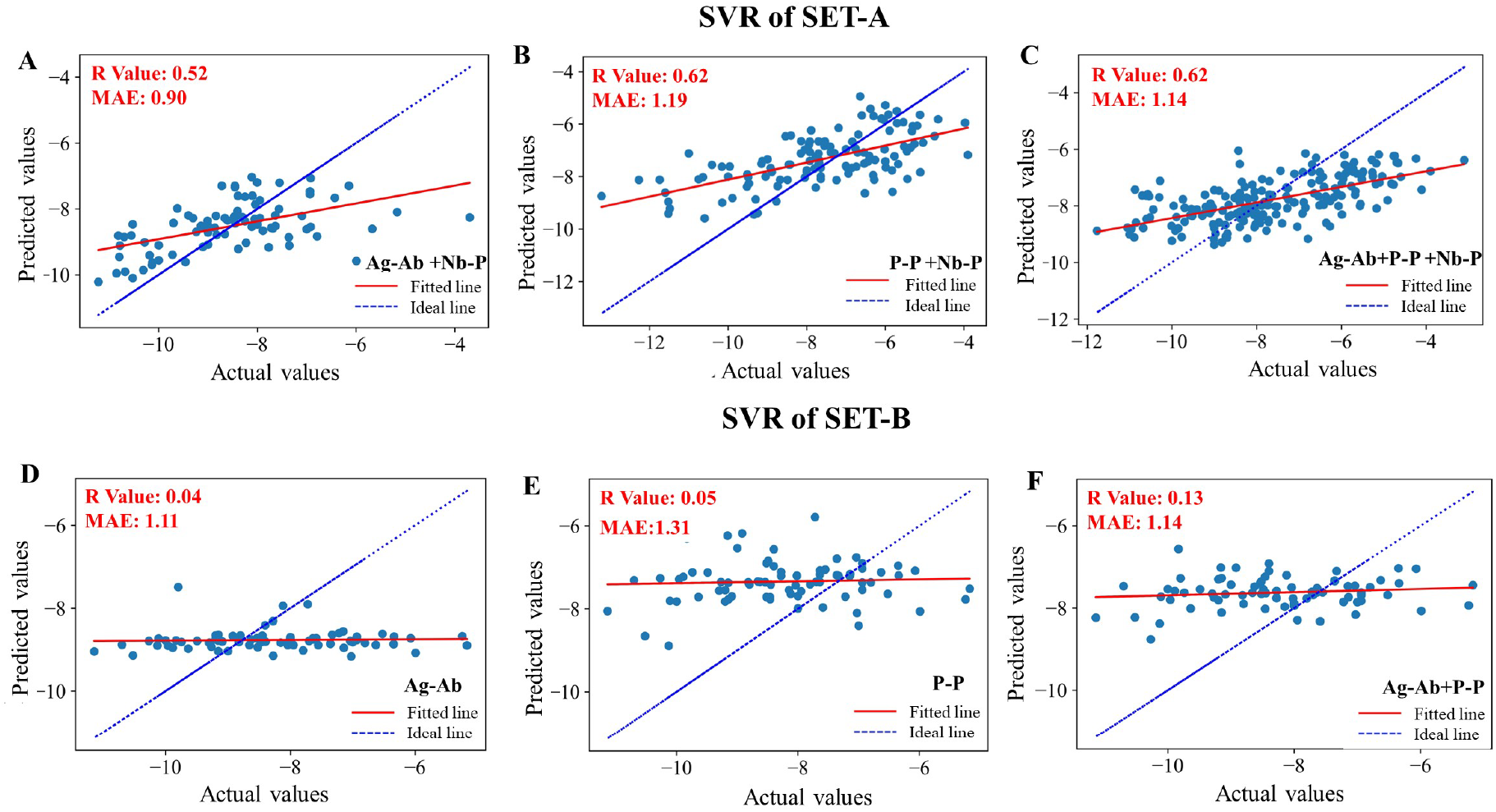
Comparison of *K*_*d*_ values predicted by SVR models for the test data against the actual experimental values, when only uncorrelated features obtained through NETCORE are considered for set A (top panel) and set B (bottom panel) data sampling schemes (Figure 3) with the training data comprising: A) 80% data of both Ag-Ab and Nb-P complexes data; B) 80% data of P-P and Nb-P complexes; C) 80% data of all protein complexes; D) all Ag-Ab complexes; E) all P-P complexes; F) all of Ag-Ab and P-P complexes. The test data for (A), (B) and (C) are the remaining 20% data and that for (D), (E) and (F) are 76 Nb-P complexes. Blue dashed line corresponds to the diagonal y = x line and red line is the linear fit to the data points representing the extent of correlation between the predicted and the actual values.

Since nanobody is a special class of antibody, we first developed a SVR model with the data on Ag-Ab complexes and Nb-P complexes with classic 80:20 split for training vs testing purposes. In Figure 6, we can notice that the R value of the predicted vs the actual values is only 0.52 and the MAE is about 0.90. Instead, when we considered P-P and Nb-P complexes for training, our performance improved slightly resulting with a R value of 0.62 but MAE increased to 1.19. On the other hand, when we included all the protein complexes, with 80:20 for training:testing, the R value remained at 0.61 but MAE slightly decreased to 1.14. It is clear that increase in number of data by including both Ag-Ab and P-P complexes, didn’t improve the performance, drastically.

When we trained the SVR model, with Ag-Ab and/or P-P complexes *i*.*e*., excluding the Nb-P complexes during training and tested the model on Nb-P complexes, the performance is extremely bad (Figure 6D–F) with a R value less than 0.14 and MAE greater than 1.04. The performance of the SVR, although, not great it is better than the predictability of popular servers when the model was trained with Nb-P complexes, but the performance is comparable to them when Nb-P complexes were excluded from the training.

### 3.4 Improved Performance with Random Forest Model

In order to improve the predictive capability, we chose Random Forest (RF), an ensemble based ML model, that builds decision trees from the subset of features and data. Initially we employed recursive feature elimination technique on the uncorrelated/weakly correlated list of features obtained from NETCORE algorithm to determine top 20 important features. Further, out of these 20 features, we explored various possible combinations of 10 important features to identify the best-performing model. During exploration, we did not fix any random state, thereby, allowing for multiple iterations to capture the best 10 features across different runs. There are two random states in a random forest model – one for splitting up of data for training vs testing and the other random state defines the important features and the decision trees of the model. The best random states were determined sequentially through a two stage process each involving 50,000 iterations; the best data split is determined in the first stage followed by the best model random state in the second stage. Figure S3 shows the histogram of R-value for the second stage of 50,000 iterations and the random state corresponding to the highest R-value is identified as the “cherrypicked” model. The average and highest R values observed during the exploration are listed in Table 2. This iterative approach ensures robust selection of the model.

**Table 2:**
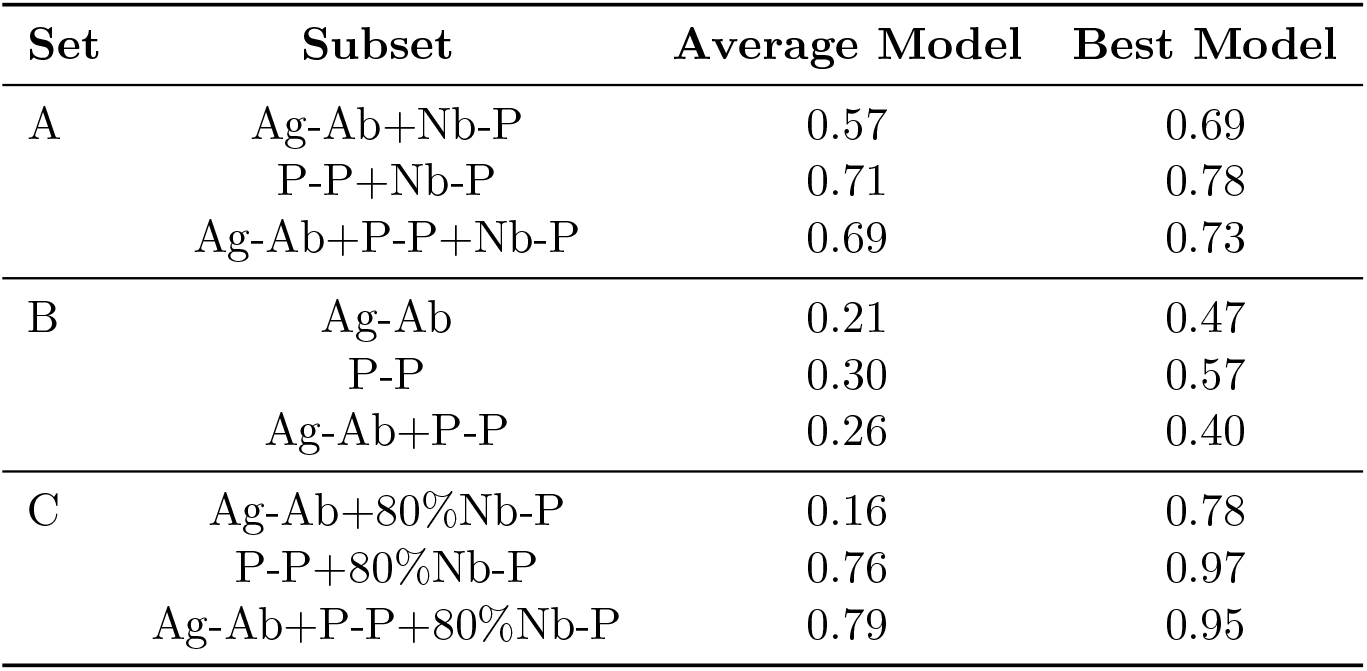
Pearson Correlation Coefficient of Average and the Best Random Forest Models.

| Set | Subset | Average Model | Best Model |
| --- | --- | --- | --- |
| A | Ag-Ab+Nb-P | 0.57 | 0.69 |
|  | P-P+Nb-P | 0.71 | 0.78 |
|  | Ag-Ab+P-P+Nb-P | 0.69 | 0.73 |
| B | Ag-Ab | 0.21 | 0.47 |
|  | P-P | 0.30 | 0.57 |
|  | Ag-Ab+P-P | 0.26 | 0.40 |
| C | Ag-Ab+80%Nb-P | 0.16 | 0.78 |
|  | P-P+80%Nb-P | 0.76 | 0.97 |
|  | Ag-Ab+P-P+80%Nb-P | 0.79 | 0.95 |

Figure 7 demonstrates the performance of the best *i*.*e*., the cherry-picked model for each dataset as per different data sampling schemes discussed in Methods section. The features corresponding to the cherry-picked RF model for each of the data sampling scheme *i*.*e*., for the each of the plots in Figure 7 is listed in Table S3. Some of the common features across these RF models are the number of polar uncharge amino acids of receptor or that in ligand, percentage of apolar residues in non-interacting surface, distance dependent statistical potentials, and quasi-chemical potential derived from the interfacial regions of protein complex. Overall, the performance of RF model is better than the SVR model (Figure 6) for the corresponding dataset.

**Figure 7:**
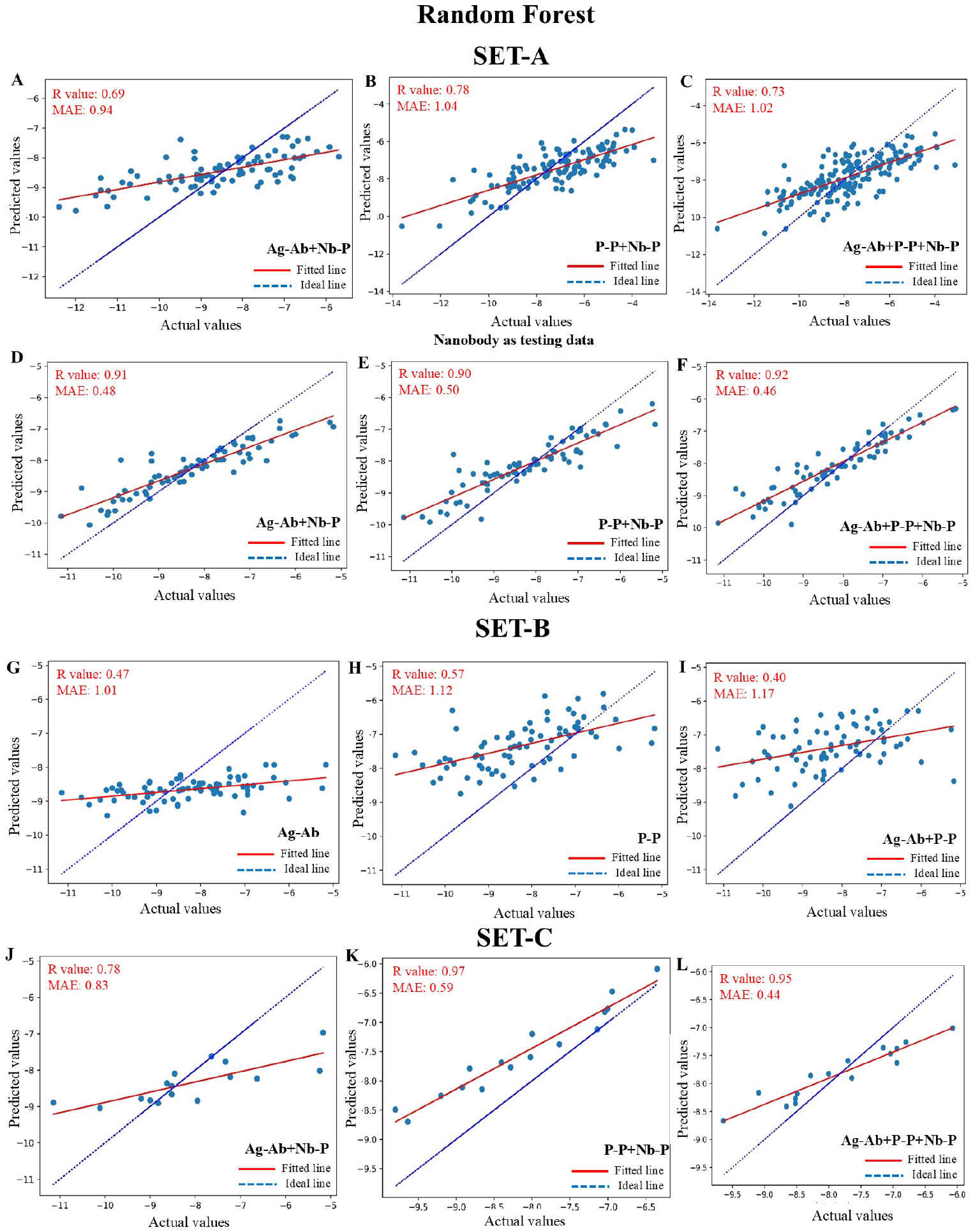
Evaluation of the performance of the cherry-picked RF model: (A),(B), and (C) are models trained on 80% data and tested on 20% data of set A *i*.*e*., Ag-Ab+Nb-P; P-P+Nb-P and Ag-Ab+P-P+Nb-P complexes, respectively; (D),(E), and (F) are models as in (A),(B) &(C) but tested exclusively on all Nb-P complexes; (G),(H), and (I) are models trained on Ag-Ab; P-P and Ag-Ab+P-P complexes, respectively, and tested on Nb-P complexes (set B); (J),(K), and (L)are models trained on Ag-Ab+80%Nb-P; P-P+80%Nb-P and Ag-Ab+P-P+80%Nb-P complexes, tested on unseen 20% Nb-P data (set C). Blue dashed line corresponds to the diagonal y = x line and red line is the linear fit to the data points representing the extent of correlation between the predicted and the actual values.

In set A sampling scheme with 80:20 data split for training and testing purposes, the R value for the Ag-Ab+Nb-P complexes dataset improved to 0.69 in RF model compared to 0.52 in SVR Model. Similarly, the R values for P-P+Nb-P and Ag-Ab+P-P+Nb-P complexes are 0.78 and 0.73, respectively, in contrast to the R value of 0.62 in Figures 6B–C. Here again, we observe that the performance of the RF model is good when the model is trained with P-P+Nb-P complexes data, inclusion of Ag-Ab data or considering Ag-Ab+Nb-P complexes resulted in slightly poor performance. When we tested the RF model obtained for the three data subsets of set A data, exclusively on all Nb-P complexes, the performance of all the models are good with R values in the range 0.90–0.92 and their MAE in the range 0.46–0.50. Interestingly, the model obtained by training on 80% of Ag-Ab+P-P+Nb-P complexes outperformed slightly compared to the performance of the Ag-Ab+Nb-P and P-P+Nb-P dataset models.

We should realize that in the Figures 7D–F, the respective RF models had already seen 80% of the tested Nb-P data. Hence, we wanted to test the performance of the RF model trained on data not including Nb-P complexes *i*.*e*., on either Ag-Ab or P-P or Ag-Ab+P-P (set B) datasets on predicting log *K*_*d*_ value of Nb-P complexes. In Figures 7G–I, we can notice the performance of RF on set B is only 0.47, 0.57 and 0.4, yet an improvement compared to SVR models with R values of 0.04, 0.05 and 0.13, respectively. This clearly suggests that inclusion of some Nb-P complexes during training is required for good performance. We, therefore, evaluated the predictive performance of nanobody *K*_*d*_ values by incorporating partial Nb-P data in training (80% of the data) and testing on the completely unseen remaining 20% data of Nb-P complexes (set C), however, we retained the same set of important features as in set B.

Figures 7K–L depict the excellent performance for set C data with R value of 0.97 and 0.95 for models trained on P-P+80%Nb-P and Ag-Ab+P-P+80%Nb-P data, respectively. As discussed earlier, the performance of the model (Figure 7J) trained on Ag-Ab+80%Nb-P data is lower, R=0.78, than the other two datasets. Since the MAE of RF model depicted in Figure 7L is the lower than the model in Figure 7K by 0.15 units though its R value is 0.02 less than the latter model, we will consider the former as the best ML model to predict the binding affinity of Nb-P complexes. The robustness of the model was further verified by bootstrap sampling on the test data, the orange histograms in figure S4 of the R-values obtained during 10,000 iterations, the average R value of the chosen best model is 0.95 *±* 0.02.

Finally, we tried to evaluate how much Nb-P data needs to be included for training the RF model for reasonable prediction of binding affinity of Nb-P complexes. We used the same important features as employed for the dataset C3 (Figure 7L), but varied the percentage of Nb-P data considered for training the RF model from 10% data to 80% data (dataset C3). The predictive performance of these RF model are shown in terms of R and MAE in Figure 8. The detailed performance plots along with histograms R value obtained during model random state selection and bootstrap sampling on test data are shown in figure S5.

**Figure 8:**
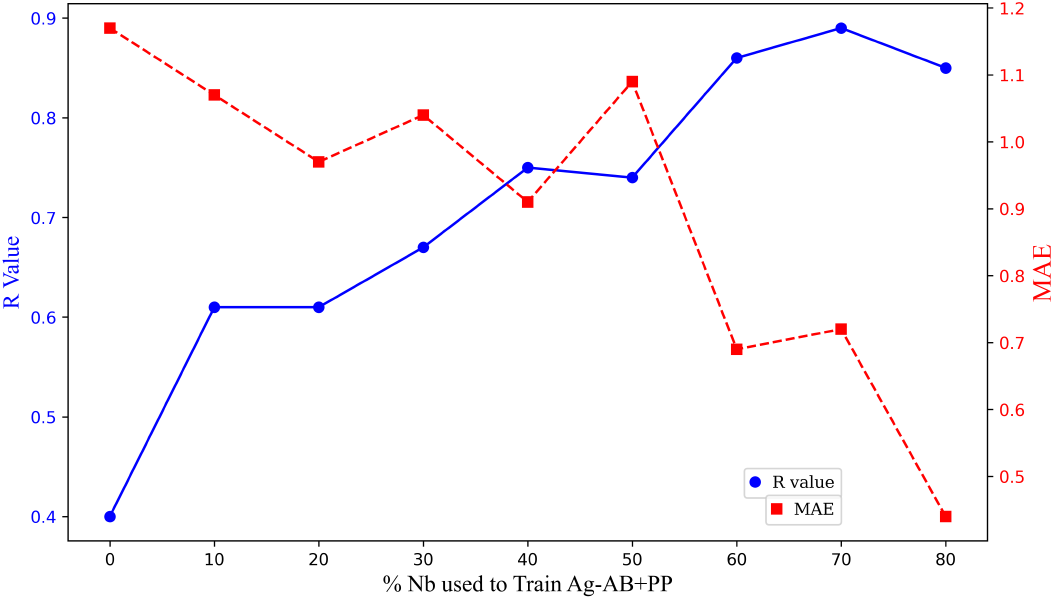
Comparing the predictive performance of RF model in terms of R and MAE values upon incremental inclusion of the Nb-P data in training the RF model

We can observe that when Nb-P data was excluded in training (dataset B3), the R value is only 0.4, with inclusion of 10% Nb-P data in training the R value drastically increases to 0.6. With further increase in percentage of data for training, we can observe either an increase in R value and/or a decrease in MAE. We can notice that inclusion of 60% Nb-P data in training results in a reasonable predictive capability with R of 0.86 and MAE of 0.69. This exercise highlights the model’s robustness even with relatively limited training data. The availability of more Nb-P data in future, can definitely help in improving the performance of the model even further. The benchmark performance with R value of 0.95 at 80% Nb-P data for training, is a clear indication of strong predictive capability of the model.

## 4 Conclusion

A computational intervention for design of nanobodies would definitely help in accelerating the development of nanobody that could target specific protein or antigen for either diagnostic or therapeutic applications. A quick estimation of binding affinity of nanobody-protein complex, therefore, is an essential requirement. Although, multiple advanced models are available to predict protein-protein interactions, the currently available popular web servers are unable to provide an accurate estimation for nanobody-protein complexes. In this work, we have cherry picked a random forest based ML model that can predict the binding affinity of nanobody-protein complex in terms of log_10_ *K*_*d*_ with a Pearson correlation coefficient of 0.95 and mean absolute error of 0.44. We found the random forest models always outperformed the support vector regression models for various datasets. We had also realized that model performance is appreciative only when we include the experimental data of some nanobody-protein complex during training.

We incorporated all the possible features typically employed in many protein-protein interaction models and finally nail down to about 10 most important features in two stages – a) elimination of highly correlated using NETCORE algorithm and b) recursive feature elimination through random forest technique. Although, nanobody is a subclass of antibody, training the model only with the data pertaining antigen-antibody complexes exhibited a sub-standard performance. The performance was consistently better when we included monomeric-protein complexes or considered only monomeric-protein complexes. This, we believe, is because both the nanobody-protein complexes and monomeric protein complexes involve only a single interface, whereas the interface is multiple and diffused in antigen-antibody complexes. Of course, the availability of more data, especially with extremely high and low *K*_*d*_ values in monomeric-protein complexes could also attribute to the better performance of the model trained with monomeric protein complexes.

## Acknowledgements

This work was partially supported by the grant MTR/2023/000699 from the Science & Engineering Research Board (SERB), India. P.M. acknowledges the financial support through Prime Minister Research Fellow Scheme, Ministry of Education, Government of India.

