## Supplementary Figures and Tables for "NanoBEP – A Machine Learning Based Tool for Nanobody Binding Energy Prediction"

Table S1: List of 134 features along with their descriptions

| Node | Feature Name | Description |
|---|---|---|
| 1 | Temperature | Temperature at which the binding affinity is measured |
| 2 | Number of charged-charged contacts | Charged: E, D, K, R. |
| 3 | Number of charged-polar contacts | Charged: E, D, K, R; polar: C, H, N, Q, S, T, Y, W |
| 4 | Number of charged-apolar contacts | Charged: E, D, K, R; apolar: A, F, G, I, V, M, P |
| 5 | Number of polar-polar contacts | Polar: C, H, N, Q, S, T, Y, W |
| 6 | Number of apolar-polar contacts | Apolar: A, F, G, I, V, M, P; Polar: C, H, N, Q, S, T, Y, W |
| 7 | Number of apolar-apolar contacts | Apolar: A, F, G, I, V, M, P |
| 8 | Percentage of apolar NIS residues (%) | Apolar: A, F, G, I, V, M, P |
| 9 | Percentage of charged NIS residues (%) | Charged: E, D, K, R. |
| 10 | Basic amino acids of receptor | Number of Basic amino acids of receptor is Antibody in the case of Ag-Ab; Nanobody in Nb-P and one of the proteins in PP complex. |

| Node | Feature Name | Description |
|---|---|---|
| 11 | Non-polar (hydrophobic) amino acids of receptor | Number of Non-polar amino acids of receptor is Antibody in the case of Ag-Ab; Nanobody in Nb-P and one of the proteins in PP complex. |
| 12 | Polar, uncharged amino acids of receptor | Number of Polar, uncharged amino acids of receptor is Antibody in the case of Ag-Ab; Nanobody in Nb-P and one of the proteins in PP complex. |
| 13 | Acidic amino acids of receptor | Number of Acidic amino acids of receptor is Antibody in the case of Ag-Ab; Nanobody in Nb-P and one of the proteins in PP complex. |
| 14 | Basic amino acids of ligand | NUmber of Basic amino acids of receptor is Antigen in the case of Ag-Ab; Protein in Nb-P and one of the proteins in PP complex. |
| 15 | Non-polar (hydrophobic) amino acids of ligand | Number of Non-polar amino acids of receptor is Antigen in the case of Ag-Ab; Protein in Nb-P and one of the proteins in PP complex. |
| 16 | Polar, uncharged amino acids of ligand | NUmber of Polar, uncharged amino acids of receptor is Antigen in the case of Ag-Ab; Protein in Nb-P and one of the proteins in PP complex. |
| 17 | Acidic amino acids of ligand | Number of Acidic amino acids of receptor is Antigen in the case of Ag-Ab; Protein in Nb-P and one of the proteins in PP complex. |
| 18 | ppdx_AGBNP | Implicit solvation energy according to the AGBNP model. |
| 19 | ppdx_ATTRACT | ATTRACT docking scoring function 32 on the structure prepared by REDUCE |
| 20 | ppdx_BSA | Total buried surface area upon binding |
| 21 | ppdx_BSA_A | Apolar buried surface area upon binding |
| 22 | ppdx_BSA_C | Charged buried surface area upon binding |
| 23 | ppdx_BSA_P | Polar buried surface area upon binding |
| 24 | ppdx_CDIE_ELEC | Electrostatic interaction as computed in a constant dielectric model |
| 25 | ppdx_CDIE_TOT | Total interaction energy as computed in a constant dielectric model |
| 26 | ppdx_CDIE_VDW | Van der Waals interaction as computed in a constant dielectric model |
| 27 | ppdx_DOPE | DOPE potential in Modeller |
| 28 | ppdx_DOPE-HR | High-resolution DOPE potential in Modeller |
| 29 | ppdx_ENM_EXP | Entropy from an Elastic Network Model where the force constant decreases exponentially with the distance |
| 30 | ppdx_ENM_R6 | Entropy from an Elastic Network Model where the force constant decreases exponentially with the distance |
| 31 | ppdx_FACTS_ASP | Apolar interaction as computed in the FACTS implicit solvent model |

| Node | Feature Name | Description |
|---|---|---|
| 32 | ppdx_FACTS_ELEC | Electrostatic interaction as computed in the FACTS implicit solvent model |
| 33 | ppdx_FACTS_GB | Generalized-Born interaction as computed in the FACTS implicit solvent model |
| 34 | ppdx_FACTS_POL | Polar interaction as computed in the FACTS implicit solvent model |
| 35 | ppdx_FACTS_TOT | Total interaction energy as computed in the FACTS implicit solvent model |
| 36 | ppdx_FACTS_VDW | Van der Waals interaction as computed in the FACTS implicit solvent model |
| 37 | ppdx_FoldX | FoldX binding score computed with the Analyse Complex tool after RepairPDB |
| 38 | ppdx_FoldX_backbone hbond | Backbone hydrogen bond component of the FoldX binding score |
| 39 | ppdx_FoldX_elec | Electrostatic component of the FoldX binding score |
| 40 | ppdx_FoldX_entropy mainchain | Main chain entropy component of the FoldX binding score |
| 41 | ppdx_FoldX_entropy sidechain | Sidechain entropy component of the FoldX binding score |
| 42 | ppdx_FoldX_sidechain hbond | Sidechain hydrogen bond component of the FoldX binding score |
| 43 | ppdx_FoldX_solvation hydrophobic | Hydrophobic solvation component of the FoldX binding score |
| 44 | ppdx_FoldX_solvation polar | Polar solvation component of the FoldX binding score |
| 45 | ppdx_FoldX_vdw | Van der Waals component of the FoldX binding score |
| 46 | ppdx_GBMV_ASP | Apolar interaction as computed in the GBMV implicit solvent model |
| 47 | ppdx_GBMV_ELEC | Electrostatic interaction as computed in the GBMV implicit solvent model |
| 48 | ppdx_GBMV_GB | Generalized-Born interaction as computed in the GBMV implicit solvent model |
| 49 | ppdx_GBMV_POL | Polar interaction as computed in the GBMV implicit solvent model |
| 50 | ppdx_GBMV_TOT | Total interaction energy as computed in the GBMV implicit solvent model |
| 51 | ppdx_GBMV_VDW | Van der Waals interaction as computed in the GBMV implicit solvent model |
| 52 | ppdx_GBSW_ASP | Apolar interaction as computed in the GBSW implicit solvent model |

| Node | Feature Name | Description |
|---|---|---|
| 53 | ppdx_GBSW_ELEC | Electrostatic interaction as computed in the GBSW implicit solvent model |
| 54 | ppdx_GBSW_GB | Generalized-Born interaction as computed in the GBSW implicit solvent model |
| 55 | ppdx_GBSW_POL | Polar interaction as computed in the GBSW implicit solvent model |
| 56 | ppdx_GBSW_TOT | Total interaction energy as computed in the GBSW implicit solvent model |
| 57 | ppdx_GBSW_VDW | Van der Waals interaction as computed in the GBSW implicit solvent model |
| 58 | ppdx_HB_BH | Number of hydrogen bonds as defined by Baker & Hubbard |
| 59 | ppdx_HB_KS | Hydrogen bond energy as defined by Kabsch & Sander |
| 60 | ppdx_HINT | Interaction energy according to the HINT model |
| 61 | ppdx_HINT_ELEC | Electrostatic interaction as computed in the HINT model |
| 62 | ppdx_HINT_VDW | Van der Waals interaction as computed in the HINT model |
| 63 | ppdx_HNC | Hydrogen network component |
| 64 | ppdx_Interfacial | Total interaction at the interface |
| 65 | ppdx_IPRO | Interface propensity index |
| 66 | ppdx_K_PSI | K score (non-binding) |
| 67 | ppdx_LIG & R - ppdx_TOTAL | Ligand and receptor total interaction score |
| 68 | ppdx_NNN | Non-nucleic nucleic interaction score |
| 69 | ppdx_POPS | Percent overlap of the protein surface |
| 70 | ppdx_RMSD | Root mean square deviation from the native structure |
| 71 | ppdx_SASA | Solvent accessible surface area |
| 72 | ppdx_VDW | Van der Waals interaction score |
| 73 | ppdx_VDO | Van der Waals (OUT) interaction score |
| 74 | ppdx_VDO_TOTAL | Van der Waals total score |
| 75 | ppdx_VALENCE | Valence score in protein-ligand complex |
| 76 | ppdx_VAN | Van der Waals score |
| 77 | ppdx_XY | XY component score |
| 78 | ppdx_XYZ | XYZ component score |
| 79 | ppdx_ZY | ZY component score |
| 80 | ppdx_ZZ | ZZ component score |
| 81 | ppdx_SOAP-PP-Pair | SOAP potential for protein-protein interfaces |
| 82 | ppdx_SOAP-Protein-OD | SOAP potential for protein structures |
| 83 | ppdx_ZRANK | ZRANK docking scoring function |
| 84 | ppdx_ZRANK2 | ZRANK2 docking scoring function (with the -R option) |
| 85 | ppdx_ipot_aace167 | AACE167 iPot contact potential |
| 86 | ppdx_ipot_aace18 | AACE18 iPot contact potential |

| Node | Feature Name | Description |
|---|---|---|
| 87 | ppdx_ipot_aace20 | AACE20 iPot contact potential |
| 88 | ppdx_ipot_rrce20 | RRCE20 iPot contact potential |
| 89 | ppdx_pyDock | PyDock docking scoring function (with pydock setup followed by pydock dockser) |
| 90 | ppdx_pyDock_desolv | Desolvation component of pyDock scoring function |
| 91 | ppdx_pyDock_elec | Electrostatic component of pyDock scoring function |
| 92 | ppdx_pyDock_vdw | Van der Waals component of pyDock scoring function |
| 93 | ppdx_sticky_avg | Average "stickiness" as defined by Levy et al. |
| 94 | ppdx_sticky_tot | Total "stickiness" as defined by Levy et al. |
| 95 | AAPP Feature 1 | Optimization-based potential derived by the modified perceptron criterion (Accession # BASU010101) |
| 96 | AAPP Feature 2 | Modified version of the Miyazawa-Jernigan transfer energy (Accession # BETM990101) |
| 97 | AAPP Feature 3 | Quasichemical statistical potential for the antiparallel orientation of interacting side groups (Accession # BONM030101) |
| 98 | AAPP Feature 4 | Quasichemical statistical potential for the intermediate orientation of interacting side groups (Accession # BONM030102) |
| 99 | AAPP Feature 5 | Quasichemical statistical potential for the parallel orientation of interacting side groups (Accession # BONM030103) |
| 100 | AAPP Feature 6 | Distances between centers of interacting side chains in the antiparallel orientation (Accession # BONM030104) |
| 101 | AAPP Feature 7 | Distances between centers of interacting side chains in the intermediate orientation (Accession # BONM030105) |
| 102 | AAPP Feature 8 | Distances between centers of interacting side chains in the parallel orientation (Accession # BONM030106) |
| 103 | AAPP Feature 9 | Distance-dependent statistical potential (only energies of contacts within 0–5 Angstroms are included) (Accession # BRYS930101) |
| 104 | AAPP Feature 10 | Quasichemical transfer energy derived from interfacial regions of protein-protein complexes (Accession # KESO980101) |
| 105 | AAPP Feature 11 | Quasichemical energy in an average protein environment derived from interfacial regions of protein-protein complexes (Accession # KESO980102) |
| 106 | AAPP Feature 12 | Statistical potential derived by the quasichemical approximation (Accession # KOLA930101) |
| 107 | AAPP Feature 13 | Modified version of the Miyazawa-Jernigan transfer energy (Accession # LIWA970101) |
| 108 | AAPP Feature 14 | Optimization-derived potential (Accession # MICC010101) |
| 109 | AAPP Feature 15 | Statistical potential derived by the maximization of the harmonic mean of Z scores (Accession # MIRL960101) |

| Node | Feature Name | Description |
|---|---|---|
| 110 | AAPP Feature 16 | Quasichemical energy of transfer of amino acids from water to the protein environment (Accession # MIYS850102) |
| 111 | AAPP Feature 17 | Quasichemical energy of interactions in an average buried environment (Accession # MIYS850103) |
| 112 | AAPP Feature 18 | Quasichemical energy of transfer of amino acids from water to the protein environment (Accession # MIYS960101) |
| 113 | AAPP Feature 19 | Quasichemical energy of interactions in an average buried environment (Accession # MIYS960102) |
| 114 | AAPP Feature 20 | Number of contacts between side chains derived from 1168 X-ray protein structures (Accession # MIYS960103) |
| 115 | AAPP Feature 21 | Quasichemical energy of transfer of amino acids from water to the protein environment (Accession # MIYS990106) |
| 116 | AAPP Feature 22 | Quasichemical energy of interactions in an average buried environment (Accession # MIYS990107) |
| 117 | AAPP Feature 23 | Quasichemical potential derived from interfacial regions of protein-protein complexes (Accession # MOOG990101) |
| 118 | AAPP Feature 24 | Distance-dependent statistical potential (contacts within 0–5 Angstroms) (Accession # SIMK990101) |
| 119 | AAPP Feature 25 | Distance-dependent statistical potential (contacts within 5–7.5 Angstroms) (Accession # SIMK990102) |
| 120 | AAPP Feature 26 | Distance-dependent statistical potential (contacts within 7.5–10 Angstroms) (Accession # SIMK990103) |
| 121 | AAPP Feature 27 | Distance-dependent statistical potential (contacts within 10–12 Angstroms) (Accession # SIMK990104) |
| 122 | AAPP Feature 28 | Distance-dependent statistical potential (contacts longer than 12 Angstroms) (Accession # SIMK990105) |
| 123 | AAPP Feature 29 | Statistical quasichemical potential with the partially composition-corrected pair scale (Accession # SKOJ000101) |
| 124 | AAPP Feature 30 | Statistical quasichemical potential with the composition-corrected pair scale (Accession # SKOJ000102) |
| 125 | AAPP Feature 31 | Statistical potential derived by the quasichemical approximation (Accession # SKOJ970101) |
| 126 | AAPP Feature 32 | Statistical contact potential derived from 25 X-ray protein structures (Accession # TANS760101) |
| 127 | AAPP Feature 33 | Number of contacts between side chains derived from 25 X-ray protein structures (Accession # TANS760102) |
| 128 | AAPP Feature 34 | Mixed quasichemical and optimization-based protein contact potential (Accession # THOP960101) |
| 129 | AAPP Feature 35 | Optimization-derived potential obtained for small set of decoys (Accession # TOBD000101) |

| Node | Feature Name | Description |
|---|---|---|
| 130 | AAPP Feature 36 | Optimization-derived potential obtained for large set of decoys (Accession # TOBD000102) |
| 131 | AAPP Feature 37 | Statistical potential derived by the maximization of the perceptron criterion (Accession # VENM980101) |
| 132 | AAPP Feature 38 | Environment-dependent residue contact energies (rows = helix, cols = helix) (Accession # ZHAC000101) |
| 133 | AAPP Feature 39 | Environment-dependent residue contact energies (rows = strand, cols = strand) (Accession # ZHAC000104) |
| 134 | AAPP Feature 40 | Environment-dependent residue contact energies (rows = coil, cols = coil) (Accession # ZHAC000106) |

AAPP features can be obtained from `https://www.genome.jp/aaindex/`
using the provided accession number

Table S2: Description of data set used for training and testing the ML models

| **Data** | **Training Data** | | **Testing Data** | | **Total** |
|---|---|---|---|---|---|
| **(sub)Set** | **Description** | **Count** | **Description** | **Count** | **Count** |
| A1 | 80%(Ag-Ab+Nb-P) | 334 | 20%(Ag-Ab+Nb-P) | 84 | 418 |
| A2 | 80%(P-P+Nb-P) | 480 | 20%(P-P+Nb-P) | 120 | 600 |
| A3 | 80%(Ag-Ab+Nb-P+P-P) | 753 | 20%(Ag-Ab+Nb-P+P-P) | 189 | 942 |
| A1' | 80%(Ag-Ab+Nb-P) | 334 | Nb-P | 76 | 410 |
| A2' | 80%(P-P+Nb-P) | 480 | Nb-P | 76 | 556 |
| A3' | 80%(Ag-Ab+Nb-P+P-P) | 753 | Nb-P | 76 | 829 |
| B1 | Ag-Ab | 342 | Nb-P | 76 | 418 |
| B2 | P-P | 524 | Nb-P | 76 | 600 |
| B3 | Ag-Ab+P-P | 866 | Nb-P | 76 | 942 |
| C1 | Ag-Ab+80%Nb-P | 402 | 20% Nb-P | 16 | 418 |
| C2 | P-P+80%Nb-P | 584 | 20% Nb-P | 16 | 600 |
| C3 | Ag-Ab+P-P+80%Nb-P | 926 | 20% Nb-P | 16 | 942 |

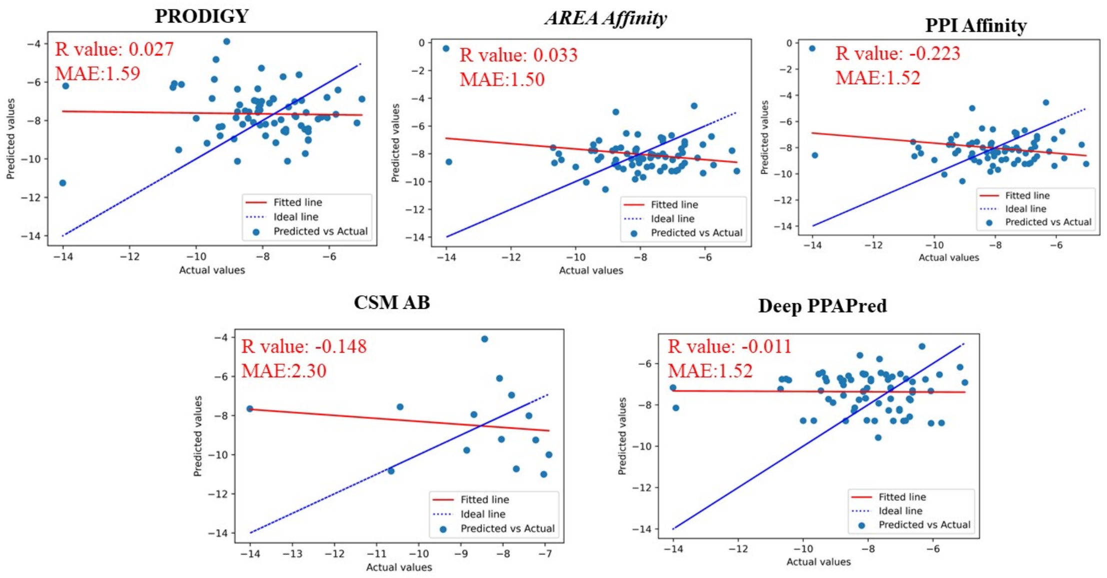


Figure S1: Comparing the performance of the popular web servers on protein-protein interactions in predicting $K_d$ values for nanobody-protein complexes

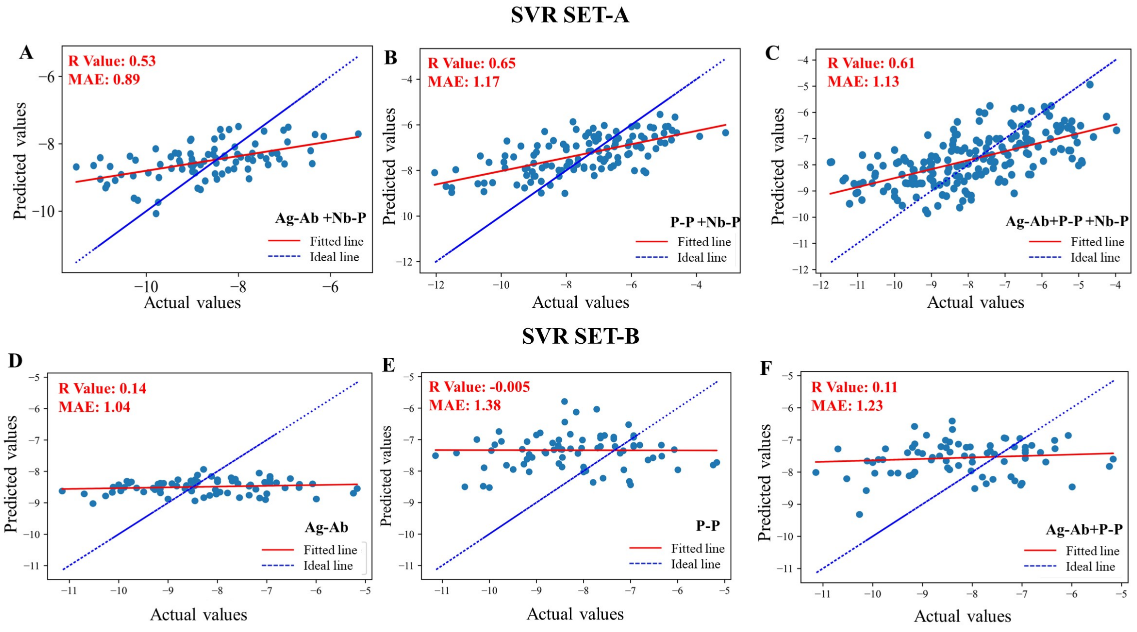


Figure S2: Comparison of $K_d$ values predicted by SVR models for the test data against the actual experimental values, when all 134 features are considered for Set A (top panel) and Set B (bottom panel) data sampling schemes (Table. S2) with the training data comprising: A) 80% data of both Ag-Ab and Nb-P complexes data (set A1); B) 80% data of P-P and Nb-P complexes (set A2); C) 80% data of all protein complexes (set A3); D) all Ag-Ab complexes (set B1); E) all P-P complexes (set B2); F) all of Ag-Ab and P-P complexes (set B3). Blue dashed line corresponds to the diagonal y = x line and red line is the linear fit to the data points representing the extent of correlation between the predicted and the actual values.

Table S3: List of top features for the *cherry-picked* random forest models obtained for different data sampling schemes

| **DataSet** | **Top Features** |
|---|---|
| A1, A1' | Temperature (1), Percentage of Charged NIS residue (9), Non-polar (hydrophobic) amino acid of receptor (11), Basic amino acids of ligand (14), Polar, uncharged amino acids of ligand (16), ppdx_AGBNP (18), ppdx_IC_TOT (67), ppdx_NIS_C (69), ppdx_NRES (71), ppdx_ipot_aace167 (85) |
| A2, A2' | Temperature (1), Percentage of apolar NIS residues (8), Polar, uncharged amino acid of receptor (12), ppdx_ENM_R6 (30), ppdx_FoldX_backbone_hbond (38), ppdx_NIS_P (70), ppdx_pyDock (89), ppdx_pyDock_vdw (92), AAPP Feature 27 (121), AAPP Feature 28 (122), AAPP Feature 37 (131) |
| A3, A3' | Temperature (1), Percentage of apolar NIS residues (8), Basic amino acids of receptor (10), Polar, uncharged amino acid of receptor (12), Acidic amino acids of receptor (13), Polar, uncharged amino acids of ligand (16), ppdx_GBMV_ELEC (47), ppdx_GBMV_TOT (50), ppdx_pyDock (89), AAPP Feature 23 (117), AAPP Feature 28 (122) |
| B1, C1 | Temperature (1), Number of charged-apolar contacts (4), Basic amino acids of receptor (10), ppdx_AGBNP (18), ppdx_FoldX_elec (39), ppdx_Rosetta_dg (78), ppdx_ipot_aace167 (85), ppdx_sticky_avg (93), AAPP Feature 21 (115), AAPP Feature 23 (117), AAPP Feature 40 (134) |
| B2, C2 | Temperature (1), ppdx_ENM_R6 (30), ppdx_GBMV_ELEC (47), ppdx_NRES (71), ppdx_Rosetta_dg (78), ppdx_ipot_aace167 (85), AAPP Feature 9 (103), AAPP Feature 14 (108), AAPP Feature 27 (121), AAPP Feature 28 (122), AAPP Feature 37 (131) |
| B3, C3 | Temperature (1), Percentage of apolar NIS residues (8), Polar, uncharged amino acid of receptor (12), Polar, uncharged amino acids of ligand (16), ppdx_Rosetta_dg (78), AAPP Feature 9 (103), AAPP Feature 23 (117), AAPP Feature 28 (122), AAPP Feature 37 (131), AAPP Feature 38 (132) |

The numbers in the brackets are the feature index as in Table S1

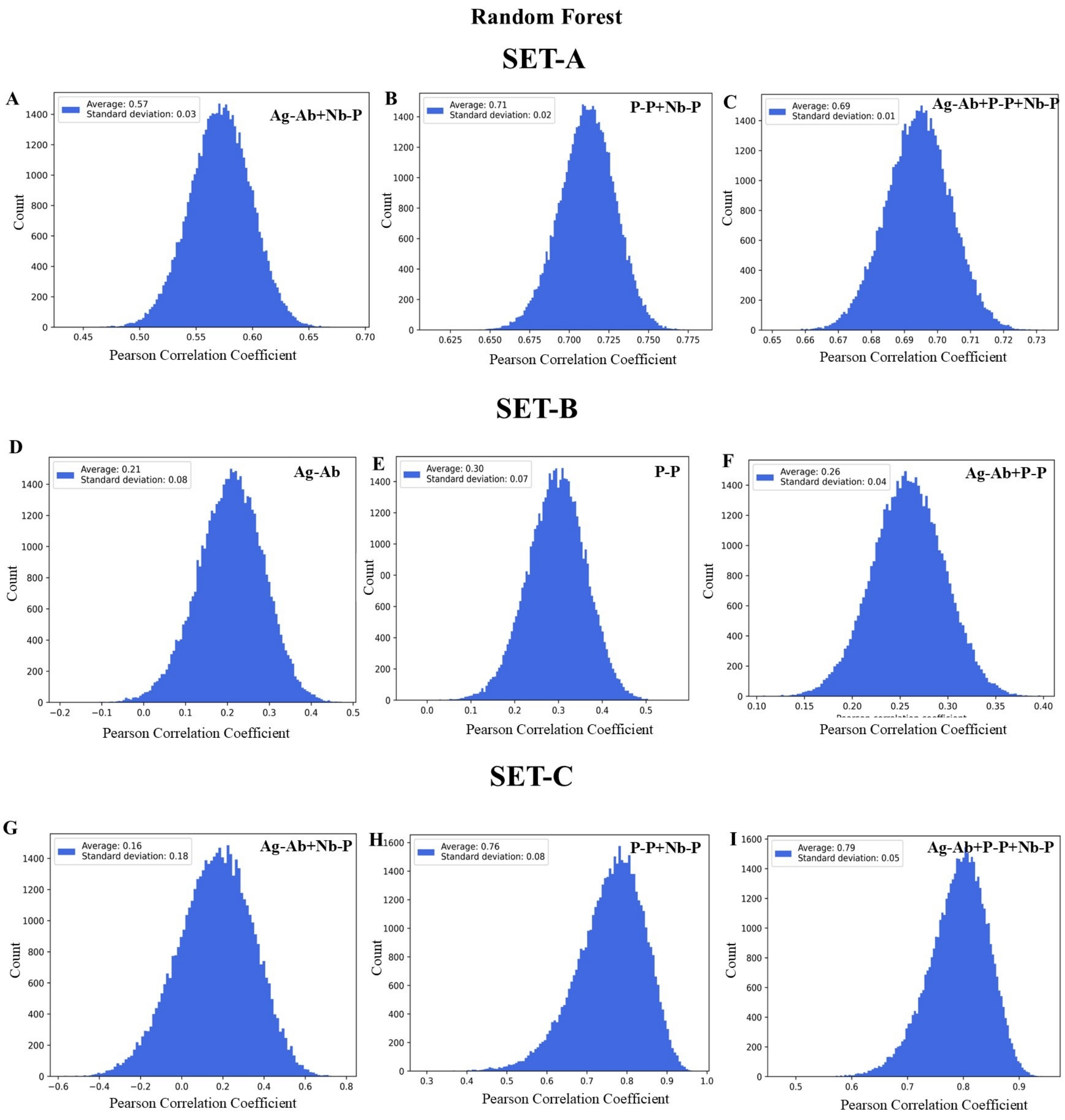


Figure S3: Histograms of Pearson correlation coefficients obtained during 50,000 iterations carried out while determining the best random forest model for set A (top panel), set B (middle panel) and set C (bottom panel) data sampling schemes. The mean and the right most models in each histogram are referred as average and the best random forest models, respectively, for the corresponding dataset.

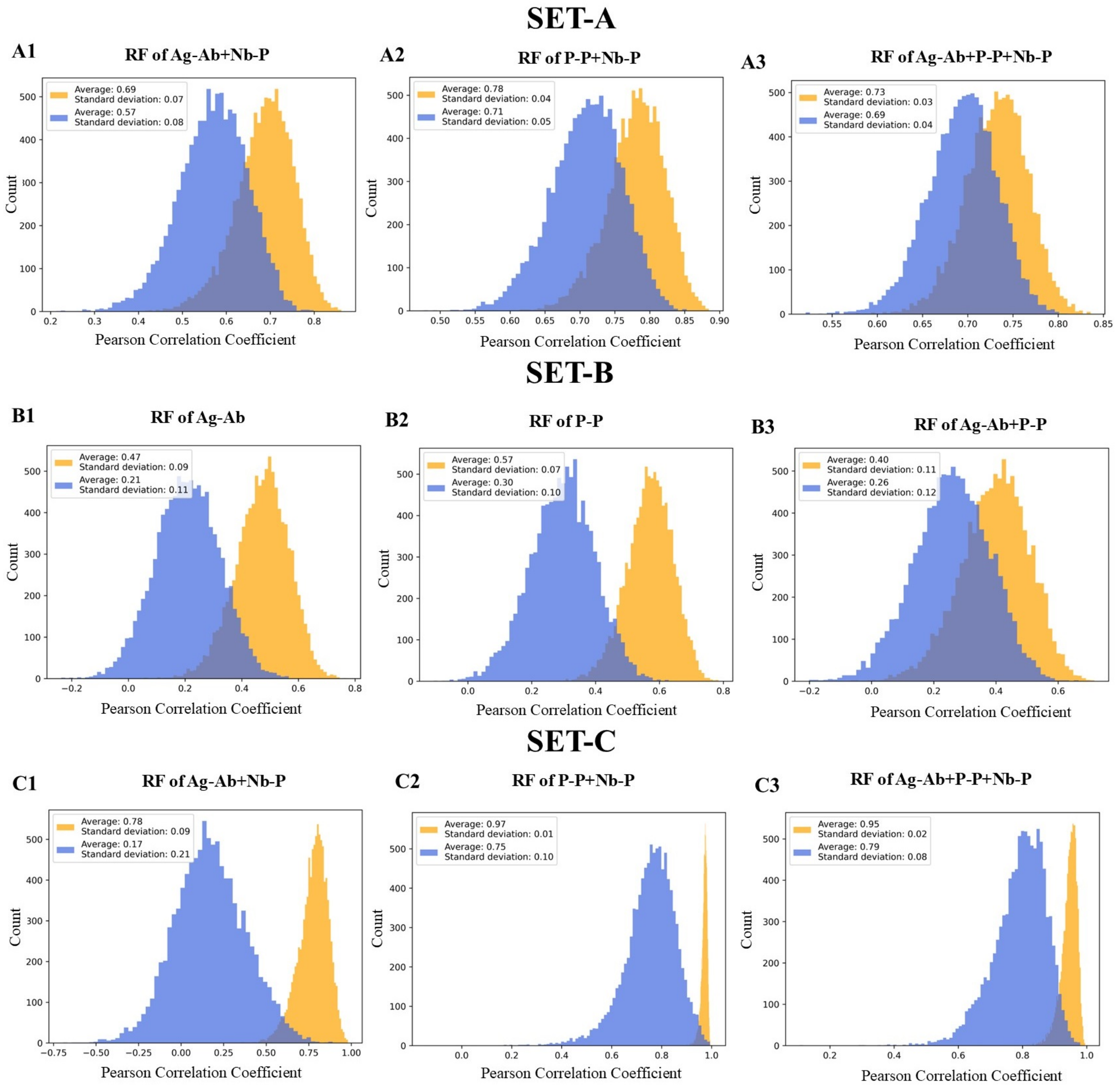


Figure S4: Histograms of Pearson correlation coefficients obtained during bootstrapping on test data while validating the robustness of the average random forest model (blue histograms) and of the best random forest model (orange histograms) identified in figure S3 for set A (top panel), set B (middle panel) and set C (bottom panel) data sampling schemes.

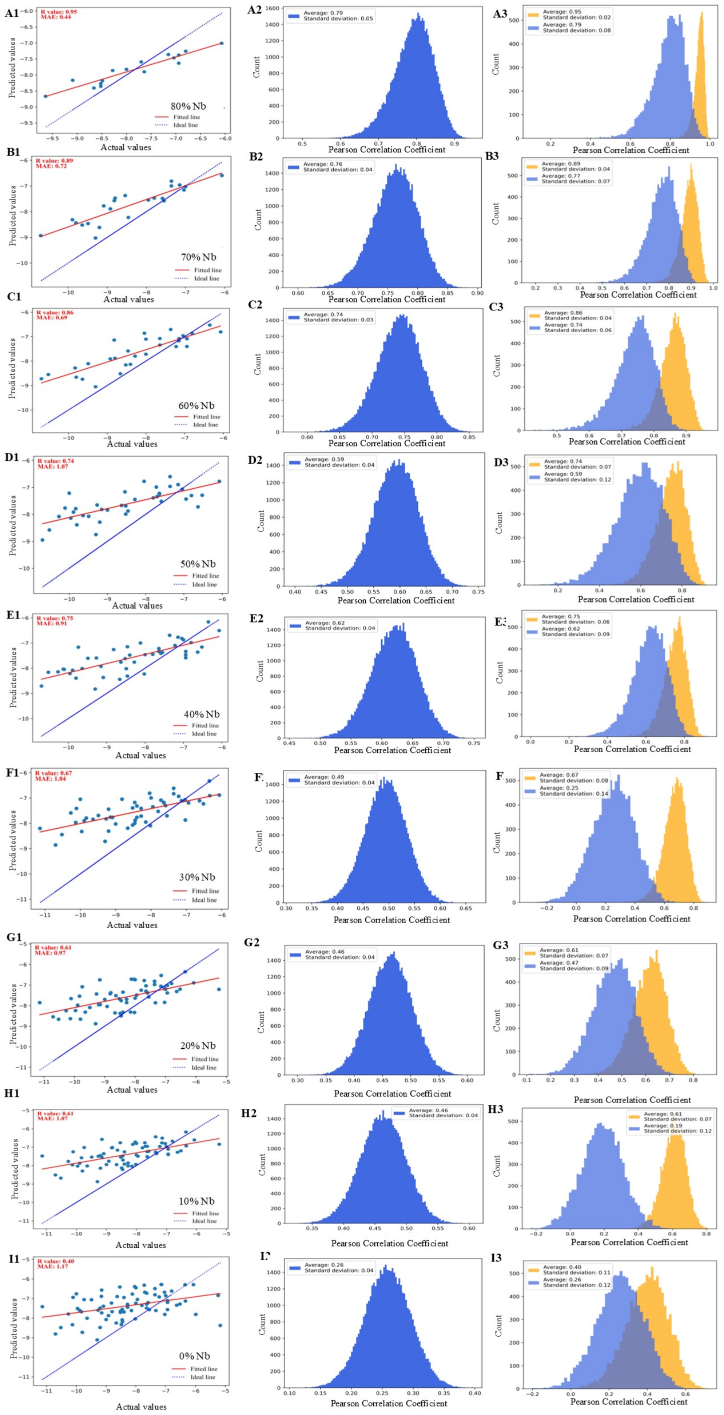

Figure S5: Performance of random forest models upon varying the percentage of Nb-P data included in training. Color code in the left, middle and right panels are as in figures S2, S3 and S4, respectively.
